# The Quantification of Drug Accumulation within Gram-Negative Bacteria

**DOI:** 10.1101/2025.05.28.656715

**Authors:** Amir George, Shivangi, Alexandra Bozan, Kendra Spencer, Austin J. Terlecky, Yong-Mo Ahn, Pamela R. Barnett, Barry N. Kreiswirth, Joel S. Freundlich

## Abstract

Intrabacterial drug accumulation, mediated by the bacterial permeability barrier, efflux, and intrabacterial drug metabolism, is of general significance to the interaction between small molecules and bacteria. For example, the ability of a small molecule to accumulate within a bacterium influences its ability to serve as a chemical probe of an intracellular protein target and/or its efficacy as an antibacterial drug discovery entity. A general method to quantitatively interrogate both intrabacterial drug accumulation and metabolism (IBDM) is presented for Gram-negative bacteria and exemplified with *Escherichia coli*, *Acinetobacter baumannii*, *Klebsiella pneumoniae*, and *Pseudomonas aeruginosa* in both single-compound and high-throughput formats. The liquid chromatography-mass spectrometry based platform does not depend on drug labelling and its utility is highlighted through the demonstrated correlation of drug accumulation with drug minimum inhibitory concentration (MIC) in both wild type and efflux deficient strains of *E. coli* and a matched pair of *K. pneumoniae* clinical and laboratory strains of varying degrees of drug resistance. Furthermore, an investigation of drug synergy implicates the selective enhancement of the accumulation of one drug by its partner therapy. Finally, a high-throughput format is validated and deployed which provides a readily adaptable approach to screening assays. We anticipate the further applications of this platform to both the translational and the fundamental studies of the interactions of small molecules with bacteria.

## INTRODUCTION

Antibacterial drug resistance represents a global health crisis.^1^ Left unchecked, annual global deaths are predicted to reach an estimated 10 million by 2050 and cost in the hundreds of trillions of dollars US.^2^ Gram-negative bacteria, in particular, represent a growing global health concern because of the emergence of multi-drug resistant (MDR) and pan-drug resistant clinical isolates.^3^ Currently, the most serious infections are nosocomial and are typically caused by *Enterobacteriaceae* spp., *Pseudomonas aeruginosa*, and *Acinetobacter* spp.^4^ In addition, community associated multi-drug resistant (MDR) infections are becoming increasingly prevalent and are most commonly associated with extended-spectrum ß-lactamase-producing *Escherichia coli* and *Neisseria gonorrhoeae*.^4^ The United States Centers for Disease Control and Prevention has classified many of these pathogens as serious threats.^3^ Notable among the urgent risks are carbapenem-resistant *Enterobacteriaceae* (CRE) which cause over 9,000 infections annually in the United States and are estimated to have a 50% mortality rate.^3,5^ Many strains possess the *K. pneumoniae* plasmid encoding a carbapenem-resistant metallo-β-lactamase to confer significant resistance to β-lactams including carbapenems, which are considered to be a treatment of last resort for MDR infections.^3,4^ In the United States alone, Gram-negative MDR pathogens have resulted in over 42,000 nosocomial infections per year, creating substantial economic strain on the healthcare system.^4^ However, in spite of the rising threat of MDR Gram-negative bacterial infections, only one new class of antibacterial (the triazaacenaphthylene gepotidacin) has been approved against these pathogens in the last fifty years.^6,7^

A major challenge in the fundamental studies of small molecule-bacteria interactions and in antibacterial chemotherapy is the consideration of the engagement of one or more intracellular protein targets by a small molecule.^8,9^ Bacteria have evolved highly effective defense mechanisms to counteract the presence of xenobiotics by limiting their accumulation within the cell itself (Fig. 1). In Gram-negative bacteria, this accumulation hurdle is especially prominent^10,11^ and the cellular features of the resistance mechanism comprise: 1) a dual-membrane envelope, consisting of a lipopolysaccharide (LPS) coated outer membrane, a thin layer of peptidoglycan, and a phospholipid bilayer inner membrane, acting as a formidable barrier to the penetration of small molecules;^12,13^ 2) an efficient multi-drug efflux pump system from the intracellular environment;^14-^16 and 3) drug-metabolizing enzymes transforming the antibacterial into nontoxic metabolite/s.^17,18^ These features constitute a primary challenge to the discovery of novel antibacterials since reduced drug accumulation leads to diminished target engagement affording attenuated whole-cell efficacy with or without an increased probability for the emergence of drug resistance. The issue of reduced drug accumulation may also be relevant to infection biology with regard to consideration of persisters.^19^

**Fig. 1.**
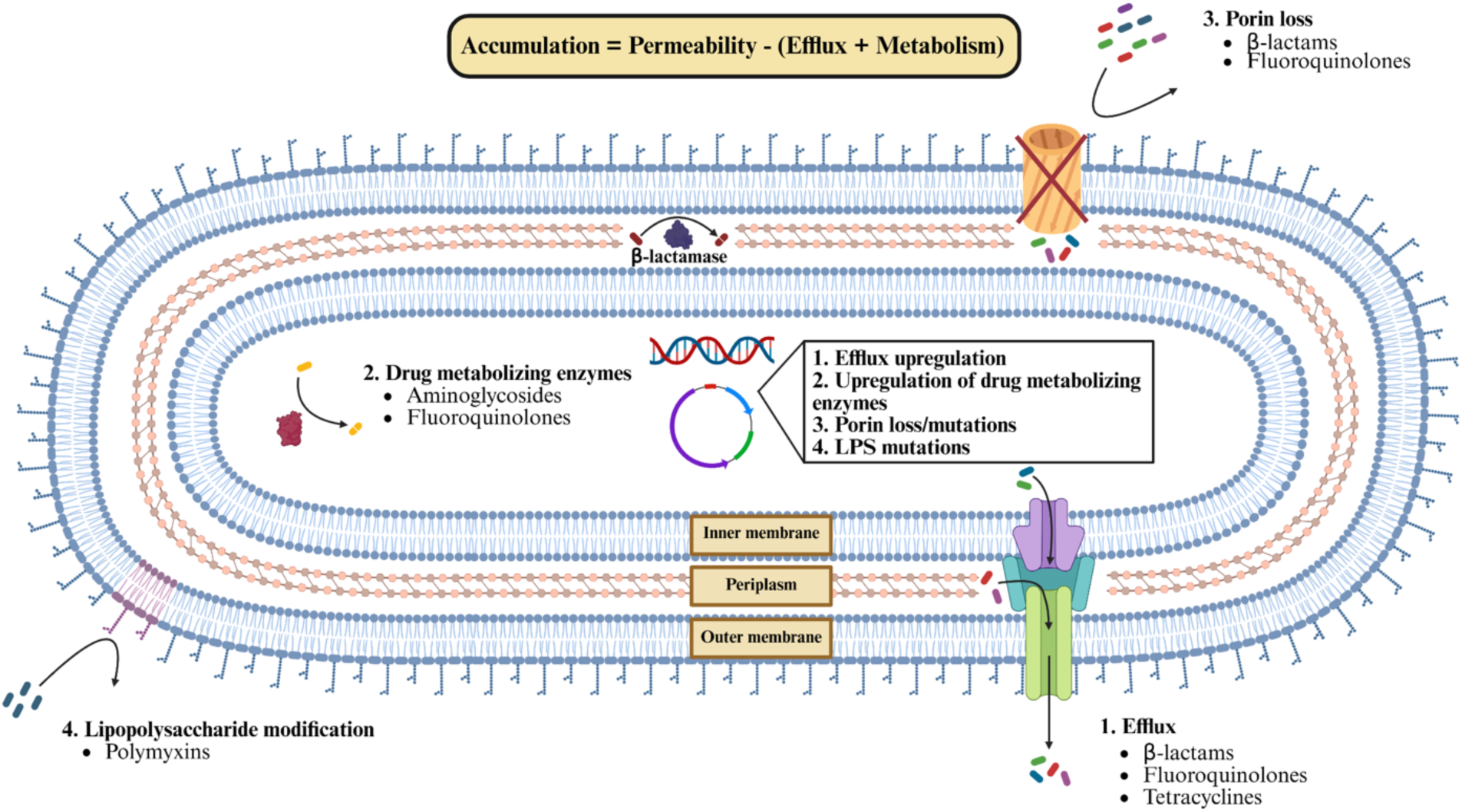
| Factors affecting intrabacterial small molecule accumulation. An overview of factors that influence intrabacterial small molecule accumulation including diffusion through porin channels, drug efflux and metabolism, and LPS modifications. Examples of drugs susceptible to these factors are listed. Created with BioRender.

In this work, we begin with the presentation of a novel intrabacterial drug accumulation and metabolism (IBDM) protocol capable of directly interrogating the dynamics of both accumulation and metabolism within a range of Gram-negative bacteria in single-compound and high-throughput (htIBDM) formats. With time- and concentration-dependent profiling, we demonstrate the utility of IBDM in characterizing the accumulation of parent drug, as well as its use in the mechanistic study of drug synergy and shedding light on the relationship between plasmid-mediated multidrug resistance and intrabacterial drug accumulation in a clinical isolate of *K. pneumoniae*.

## Results

### IBDM assay optimization with *E. coli*

We commenced with the adaptation of the IBDM assay from our previous work with *Mycobacterium tuberculosis*^20–23^ and *Staphylococcus aureus*^24^ to pursue drug accumulation measurements in the wild type *E. coli* strain MG1655. Four major experimental steps exist in the protocol: 1) bacterial growth, incubation with drug, and sample collection; 2) washing and quenching; 3) bacterial cell lysis and filtration; and 4) analysis of samples via liquid chromatography-mass spectrometry (LC/MS) (Fig. 2). To optimize the protocol, bacteria were cultured in the presence of the antibacterial rifampicin, and the effects of total culture volume (15, 25, and 125 mL) and time point (10, 30, 60, and 90 min) on intrabacterial accumulation were observed (Supplementary Figs. 1 and 2), where the time points are referenced to the addition of the drug to the bacterial culture. Rifampicin was utilized because it is a lower accumulating drug in *E. coli*^25^ and we hypothesized that its use in parameter optimization would result in a more sensitive assay. No significant differences in rifampicin accumulation between the different conditions were observed. However, the 25 mL total culture volume was chosen over the 10 mL volume because of the observation of larger, more well-formed cell pellets with the 25 mL cultures. The first sample was collected at 10 min, given our above findings and previous observations from the groups of Hergenrother^26^ and Piddock^27^.

**Fig. 2.**
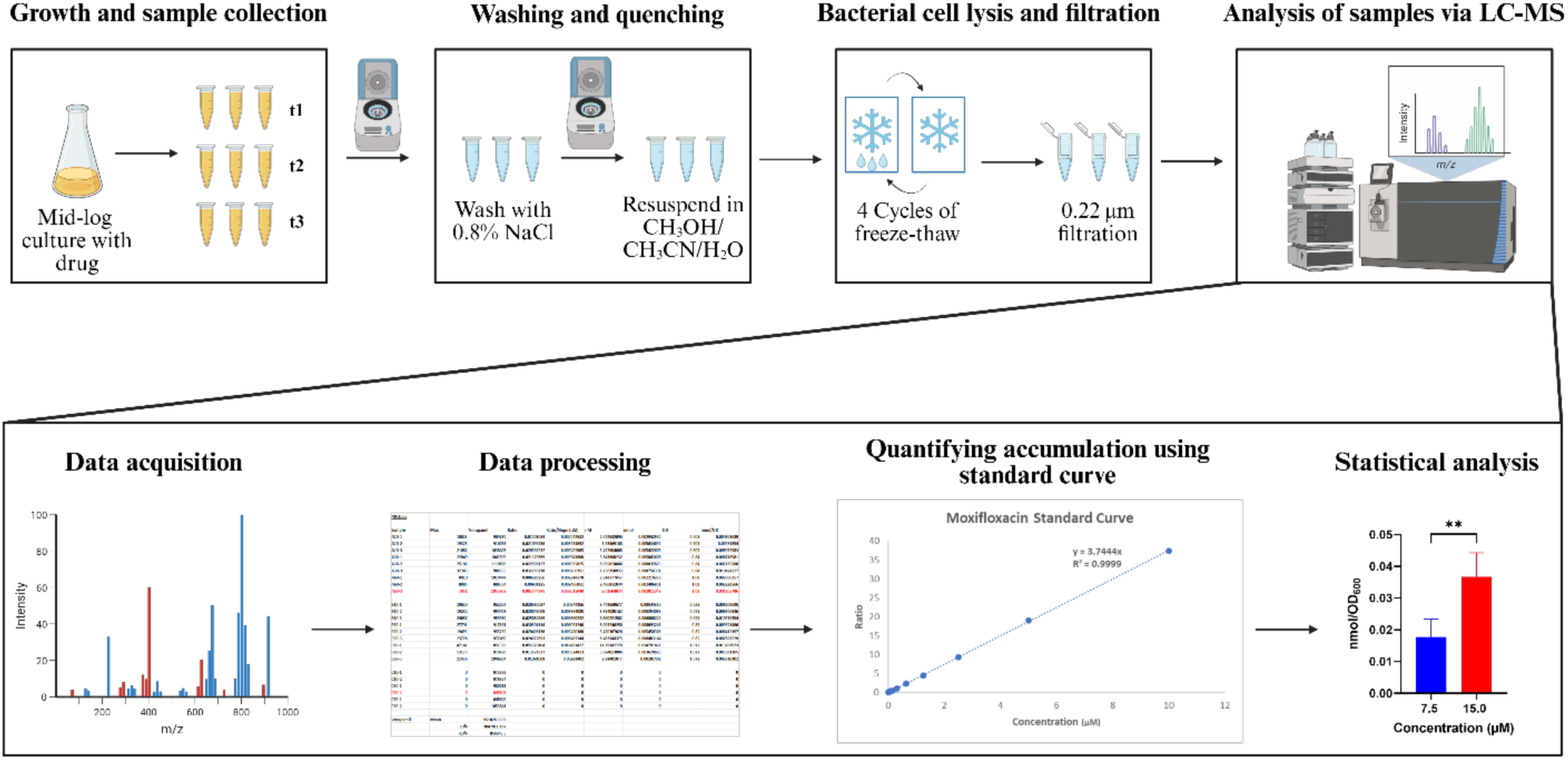
|. General IBDM assay protocol. Summary of the IBDM protocol, which allows for quantitative analysis of the accumulation of the drug and its identified metabolites within the bacteria under study. This figure was created with BioRender.

Subsequently, we sought to test the validity of using OD600 readings to normalize accumulation trends compared to colony-forming units (CFUs). We tested this by measuring the accumulation of doxycycline, as a precedented high accumulator,^26^ and rifampicin in wild type *E. coli*. No significant difference was observed between the two trends when comparing the different normalizations (Supplementary Fig. 3). We next observed changes in moxifloxacin, doxycycline, and rifampicin accumulation at the three different time points. We observed no significant time-dependent changes in intrabacterial drug accumulation at 10, 30, and 60 min at two different concentrations of each drug (Supplementary Fig. 4). Therefore, all subsequent accumulation studies were pursued using the 10 and/or 60 min time points. For example, we used the 60 min timepoint to examine the effect of drug concentration added on intrabacterial drug accumulation.

Significant differences in intrabacterial accumulation of doxycycline (7.5 versus 15 µM) and rifampicin (10 versus 20 µM) were noted with respect to the different concentrations (Supplementary Fig. 5). However, no significant accumulation difference was noted between the two moxifloxacin concentrations (2.5 and 5.0 µM), potentially due to the observed nmols/OD600 values being close to the lower limit of detection (LLOD) of the drug.

Drug accumulation of doxycycline, moxifloxacin, and rifampicin in isogenic strains of *E. coli*.

Since AcrAB-TolC is one of the major efflux pumps implicated in multi-drug resistance in Gram-negative bacteria,^28^ we quantified the accumulation of this set of drugs in an efflux deficient *ΔtolC* strain. We hypothesized intrabacterial drug accumulation would be increased in the *ΔtolC* strain for drugs which exhibit greater growth inhibition, i.e., a lower MIC, in this efflux deficient strain as compared to the wild type MG1655 strain. This was observed for both doxycycline and moxifloxacin (Fig. 3) as their respective intrabacterial accumulations were significantly higher in the *ΔtolC* strain as compared to the MG1655 strain at the 60 min time point. No significant difference in rifampicin accumulation was observed between the wild type and the *ΔtolC* strain (Fig. 3), which correlates with the modest change (twofold) in MIC for rifampicin between the strains (Table 1).

**Fig. 3.**
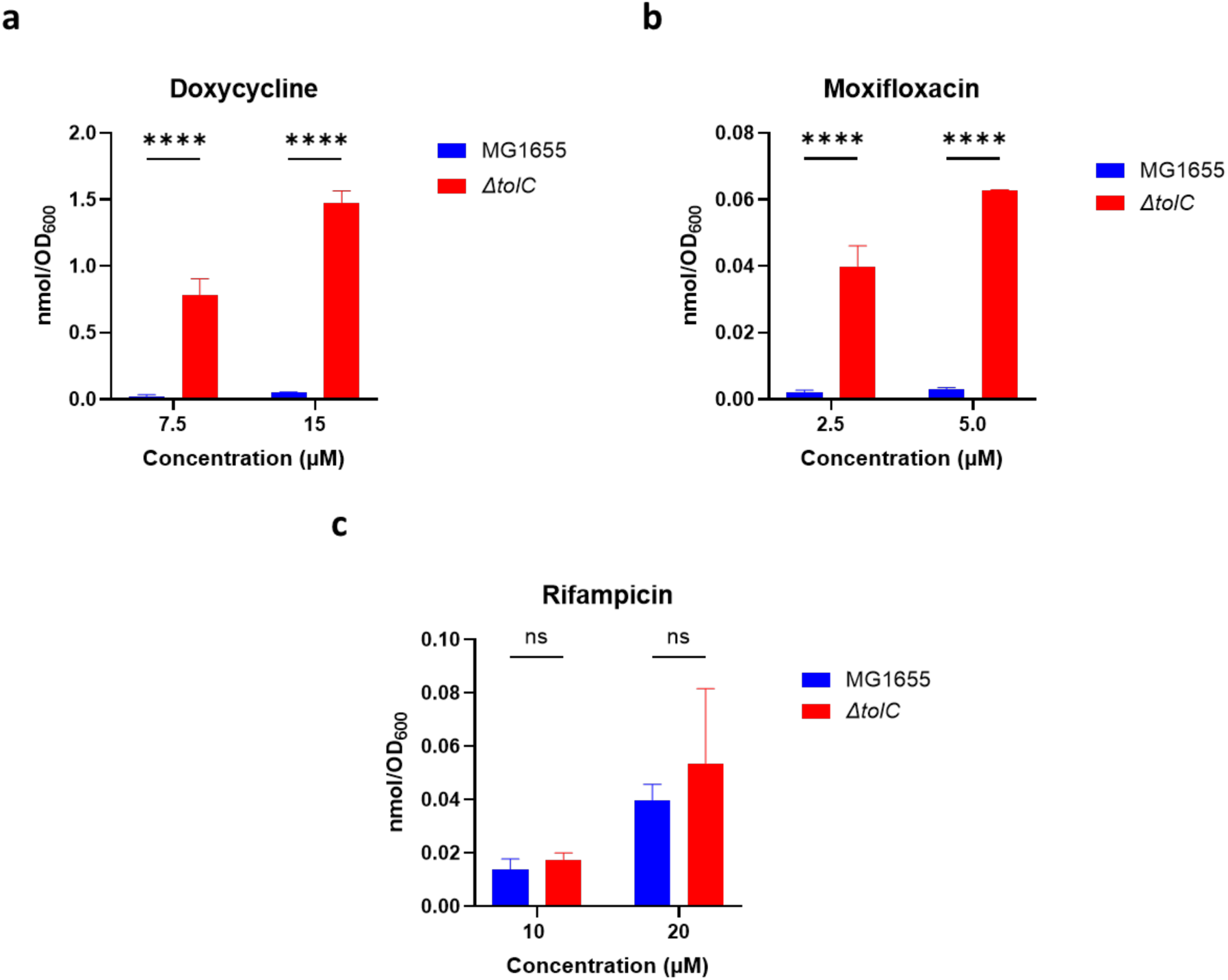
| Accumulation of moxifloxacin, doxycycline, and rifampicin in *E. coli* MG1655 and *ΔtolC* strains. a. Doxycycline at 7.5 and 15 µM, b. Moxifloxacin at 2.5 and 5.0 µM, c. Rifampicin at 10 and 20 µM. All measurements were at 60 min. Data are shown as mean ± SD for a representative of two independent experiments each conducted in quadruplicate. The amount of accumulated compound was quantified as the number of moles normalized by OD600. p-values were determined with the two-way ANOVA with Šidák post hoc test. ns p>0.05, ** p<0.01, *** p<0.001, **** p<0.0001.

**Table 1.**
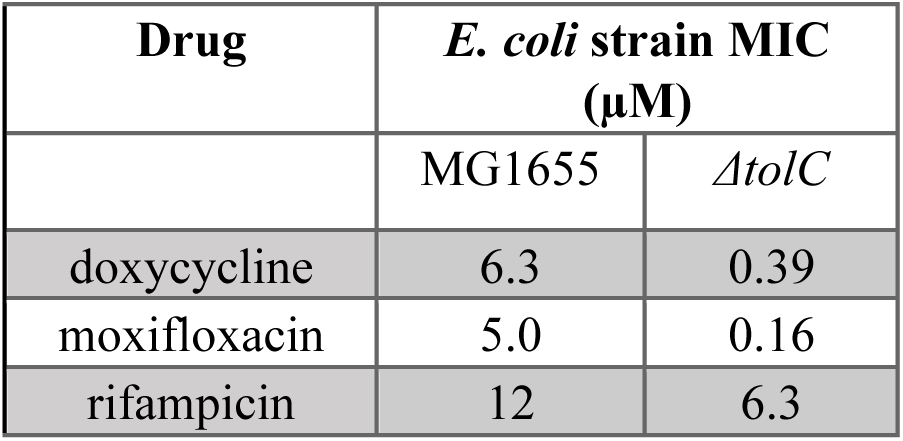
| MIC values for select antibacterial agents versus wild type and efflux-deficient *E. coli*. Each MIC value represents the average value from a minimum of two independent experiments.

| Drug | *E. coli* strain MIC (μM) |  |
| --- | --- | --- |
|  | MG1655 | *ΔtolC* |
| doxycycline | 6.3 | 0.39 |
| moxifloxacin | 5.0 | 0.16 |
| rifampicin | 12 | 6.3 |

### The IBDM platform may be extended to other Gram-negative bacteria

After leveraging the IBDM assay with *E. coli*, we sought to apply the platform to other Gram-negative bacteria of global health relevance. We selected representative strains of *Acinetobacter baumannii* (ATCC#19606)*, Pseudomonas aeruginosa* (ATCC# HER-1018), and *Klebsiella pneumoniae* (ATCC# BAA 2146) as they belong to a particularly difficult to treat group of bacteria associated with nosocomial infections known as ESKAPE bacteria.^29^ We chose rifampicin to assess against these bacteria due to its activity against the tested strains (Supplementary Table 1). The *E. coli* IBDM protocol was utilized without alteration. At a rifampicin concentration of 10 µM for all three bacteria, we observed statistically significant greater accumulation of rifampicin when incubated at a concentration of 10 µM in *A. baumannii* compared to *K. pneumoniae* and *P. aeruginosa* (Fig. 4).

**Fig. 4.**
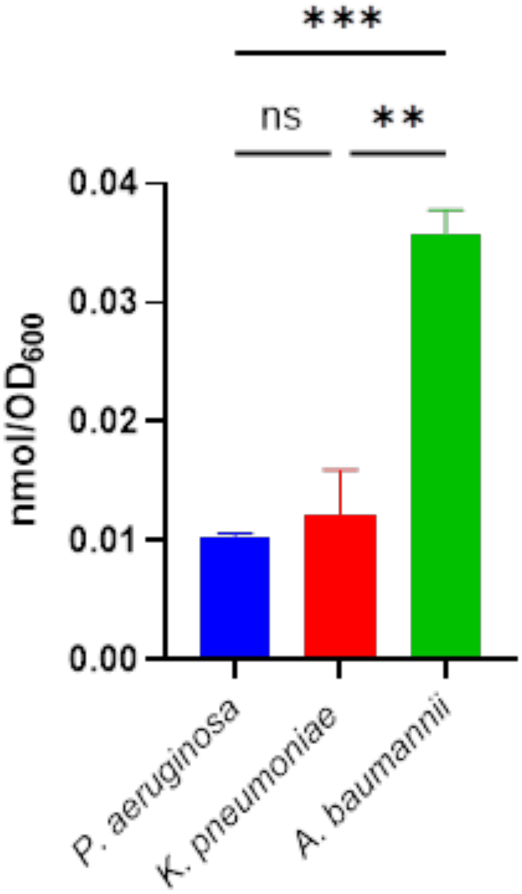
**| Rifampicin accumulation in *P. aeruginosa, K. pneumoniae, and A. baumannii.*** Rifampicin (10 µM) accumulation in *A. baumannii* (ATCC# 19606)*, P. aeruginosa* (ATCC# HER-1018), or *K. pneumoniae* (ATCC# BAA 2146) at t = 60 min. Data are shown as mean ± SD representative for a representative of two independent experiments each conducted in triplicate. The amount of accumulated compound as the number of moles was normalized by OD600. p-values were determined by one-way ANOVA with Tukey post hoc test. ns p>0.05, ** p<0.01, *** p<0.001, **** p<0.0001.

### Drug accumulation measurements may be applied to examine drug combinations

Since drug combinations can be integral to the treatment of Gram-negative MDR infections,^30^ the IBDM assay was applied to study the intrinsic drug-drug interactions. A study from the Blainey laboratory^31^ showed that indacaterol – an approved drug for the treatment of chronic obstructive pulmonary disorder^32^ – exhibited synergy with novobiocin and erythromycin against the *E. coli* MG1655 strain. Indacaterol is inactive against this *E. coli* strain.^31^ Although novobiocin is active as a single agent versus Gram-positive bacteria, it is inactive against Gram negatives presumably due to its large size and hence low permeability across the Gram-negative membrane.^25,31,33^ Therefore, we hypothesized that indacaterol increases the permeability of *E. coli* through potential perturbation of the bacterial cell membrane. Given our previous profiling of rifampicin as a relatively low accumulating drug within *E. coli* MG1655, we chose to examine its interaction with indacaterol. A checkerboard assay was performed with rifampicin and indacaterol. The fractional inhibitory concentration index (FICI)^34^ was found to be ≤0.19 (Supplementary Fig. 6), classifying the combination as synergistic (i.e., FICI ≤ 0.5). After confirming the synergistic interaction between rifampicin and indacaterol, we quantified the accumulation in the *E. coli* MG1655 strain of rifampicin alone and in the presence of indacaterol. In the presence of 50, 100, or 200 µM indacaterol, rifampicin accumulation was significantly higher than that with rifampicin alone (Fig. 5). Furthermore, no statistically significant difference was noted between the accumulation of rifampicin alone or in the presence of 25 µM indacaterol, which is lower than indacaterol’s MIC in combination.

**Fig. 5.**
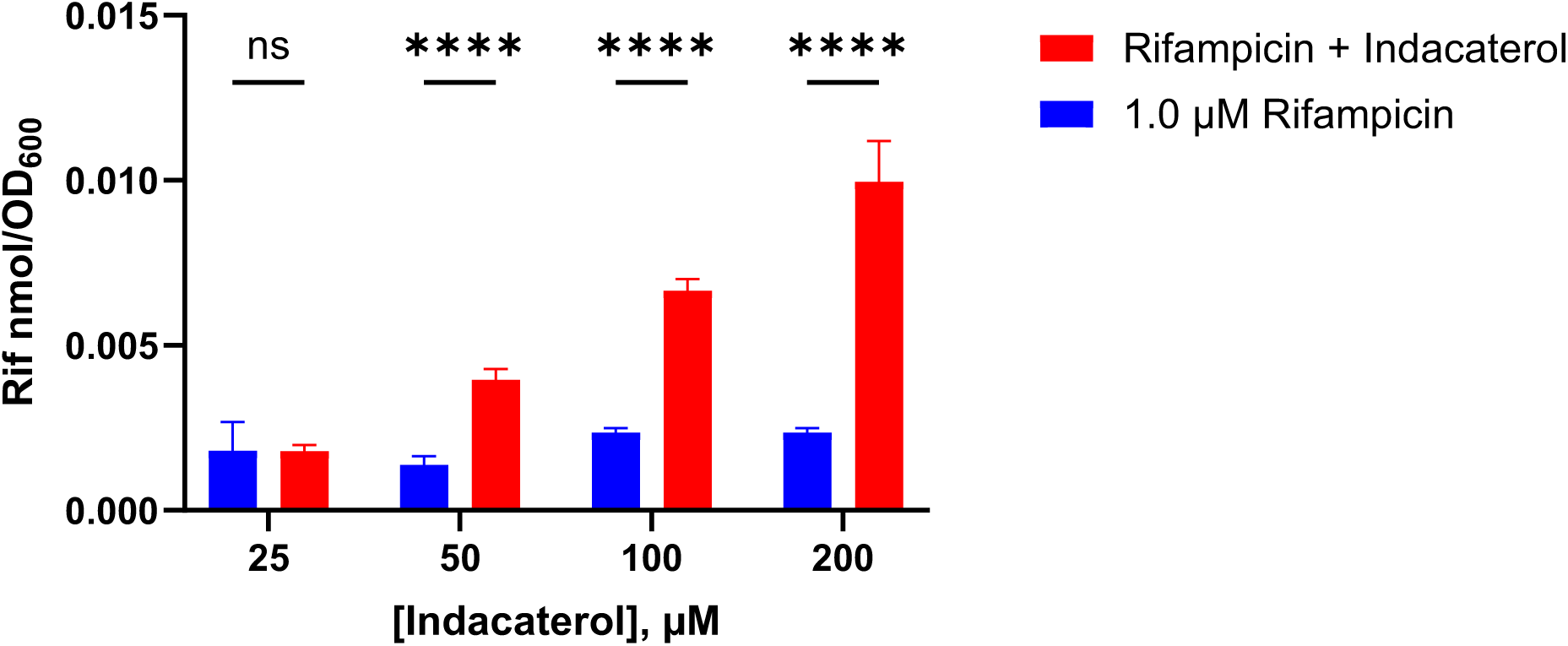
**| Rifampicin accumulation in *E. coli* MG1655 in the presence of indacaterol.** Rifampicin (1.0 µM) in the presence of varying concentrations of indacaterol (25, 50, 100 or 200 µM) was incubated with *E. coli* MG1655 for 60 min prior to measurement. Data are shown as mean ± SD representative for a representative of two independent experiments each conducted in triplicate. The amount of accumulated compound as the number of moles was normalized by OD600. p-values were determined by two-way ANOVA with Šidák post hoc test for all assays. ns p>0.05, ** p<0.01, *** p<0.001, **** p<0.0001.

To further investigate this phenomenon, we performed flow cytometry studies using the cell impermeable dye (TO-PRO-3).^35^ We observed a higher percentage of the cells taking up the dye at 200 µM indacaterol as compared to 25 µM indacaterol (Fig. 6), and these results changed little when rifampicin and indacaterol were used in combination. Furthermore, we did not observe a significant difference in dye uptake between the 1.0 µM rifampicin and DMSO control treatments (Supplementary Fig. 7), suggesting that rifampicin is not affecting cell permeability alone and that the permeabilization observed in the 1:200 rifampicin:indacaterol condition is likely mediated through indacaterol. Altogether, these results are consistent with the hypothesis that indacaterol is synergizing with rifampicin through membrane permeabilization to increase the rifampicin accumulation inside the cells.

**Fig. 6.**
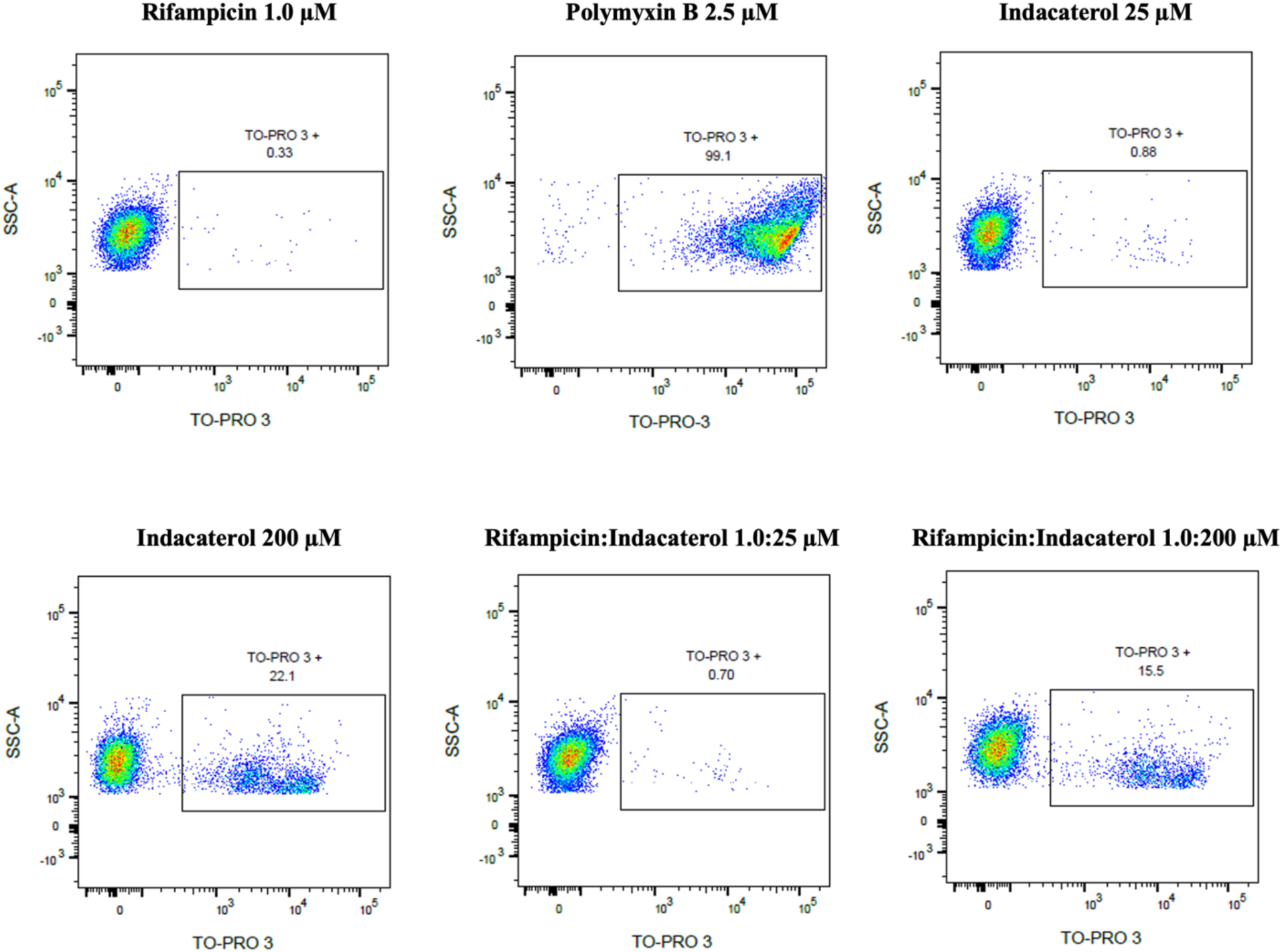
| Effect of indacaterol on *E. coli* membrane permeability to TO-PRO-3 dye measured by flow cytometry. The percent of cells that took up the dye is shown under each condition. The y-axis represents the forward scatter area and the x-axis represents the fluorescence of TO-PRO-3. Polymyxin B was used as the positive control. The results are representative of three independent experiments.

### A high-throughput platform (htIBDM) is developed

We next examined the possibility of extending the IBDM protocol to a high-throughput format (htIBDM). The protocol for htIBDM is similar to that for IBDM with a few alterations in the washing and lysis steps (Fig. 7). To validate the htIBDM, we calculated the Z′ value^36^ to be 0.70 with the *E. coli* MG1655 strain and 10 µM doxycycline. We subsequently performed a htIBDM screen with doxycycline, moxifloxacin, ciprofloxacin, rifampicin and novobiocin at concentrations of 20 and 40 µM with the *E. coli* MG1655 and *ΔtolC* strains (Fig. 8). Accumulation trends were consistent when compared with results from single compound IBDM for those same drugs (Supplementary Fig. 8). Doxycycline was the highest accumulator in both assay formats and rifampicin was the lowest accumulator. Ciprofloxacin accumulation was significantly higher than moxifloxacin accumulation in both wild type and efflux deficient *E. coli* despite both drugs belonging to the fluoroquinolone family, suggesting even seemingly minor differences in chemical structures can significantly impact accumulation. Novobiocin accumulation was lower in the htIBDM assay than in the IBDM assay, potentially due to an extra aqueous NaCl wash deemed necessary due to the comparatively higher predicted hydrophobicity of novobiocin (predicted logD = 3.4, Supplementary Table 2) leading to its proposed adherence to the bacterial membrane. Increasing the number of aqueous NaCl washes to two mitigated this problem (Supplementary Fig. 9). Overall, these results are consistent with the studies performed by Richter and colleagues.^26^

**Figure 7.**
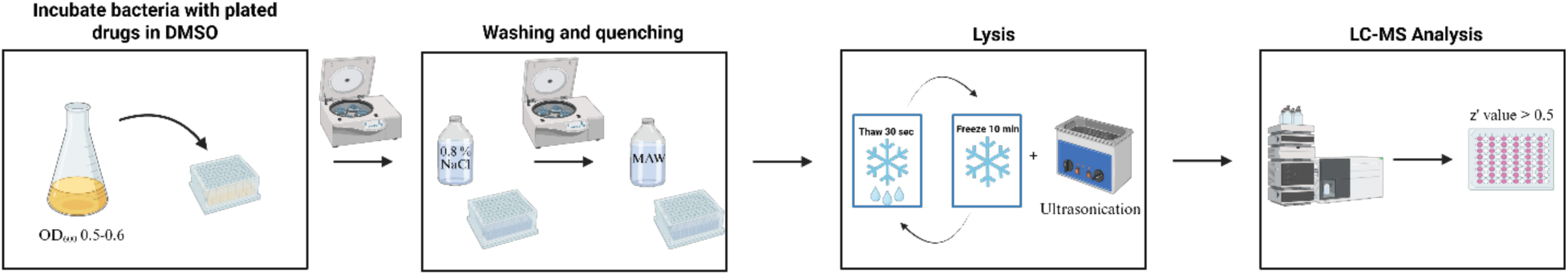
| General high-throughput IBDM (htIBDM) protocol. Summary of the htIBDM protocol which allows for quantitative analysis of drug accumulation and its identified metabolites in a 96-well plate. This figure was created with BioRender.

**Fig. 8.**
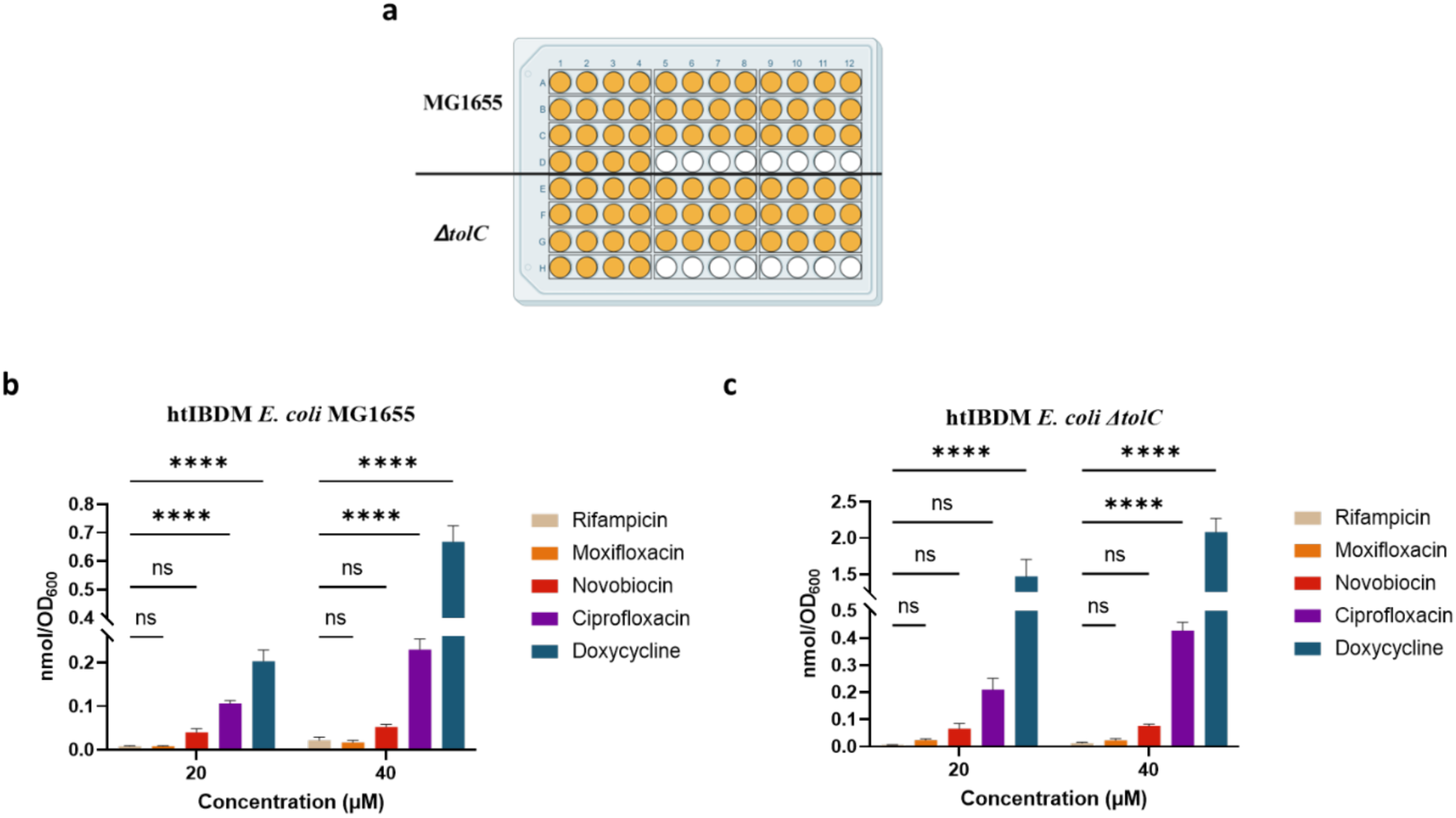
**| An exemplary htIBDM of select antibacterials with the *E. coli* MG1655 and *ΔtolC* strains.** a. Plate layout for htIBDM to measure the intrabacterial accumulation of rifampicin, moxifloxacin, novobiocin, ciprofloxacin, and doxycycline at 20 and 40 µM (yellow-orange) and DMSO treated controls (white) in quadruplicate with two *E. coli* strains. Intrabacterial accumulation of drugs in b. wild type MG1655 and c. efflux deficient *ΔtolC E. coli* strains. Data are shown as mean ± SD representative of two independent experiments each conducted in quadruplicate. The amount of accumulated compound as the number of moles was normalized by cell number as approximated by OD600. p-values were determined by two-way ANOVA with Tukey post hoc test for all assays. ns p>0.05, ** p<0.01, *** p<0.001, **** p<0.0001.

The htIBDM assay was subsequently deployed to studies with *A. baumannii* (ATCC #19606), *P. aeruginosa* (ATCC #HER-1018), and *K. pneumoniae* (ATCC #BAA 2146). The robustness of the assay was again confirmed with Z’ factors ≥ 0.5 obtained via assay validation using doxycycline (10 µM) as a positive control. We examined the intrabacterial accumulation of doxycycline, moxifloxacin, ciprofloxacin, and rifampicin at a 20 µM concentration (Fig. 9). Across all three strains and consistent with studies conducted in *E. coli* (Supplementary Fig. 8), doxycycline was the highest accumulator, rifampicin was the lowest accumulator, and ciprofloxacin accumulated higher than moxifloxacin. The accumulation of both doxycycline and ciprofloxacin was significantly lower in the *K. pneumoniae* strain used. This particular strain is a clinical isolate that exhibits multidrug resistance mediated through a combination of genes on its own chromosome and 23 plasmid-borne genes encoding drug resistance proteins.^37,38^ These observations piqued our interest in further exploring the effect of plasmid-mediated resistance on accumulation.

**Fig. 9.**
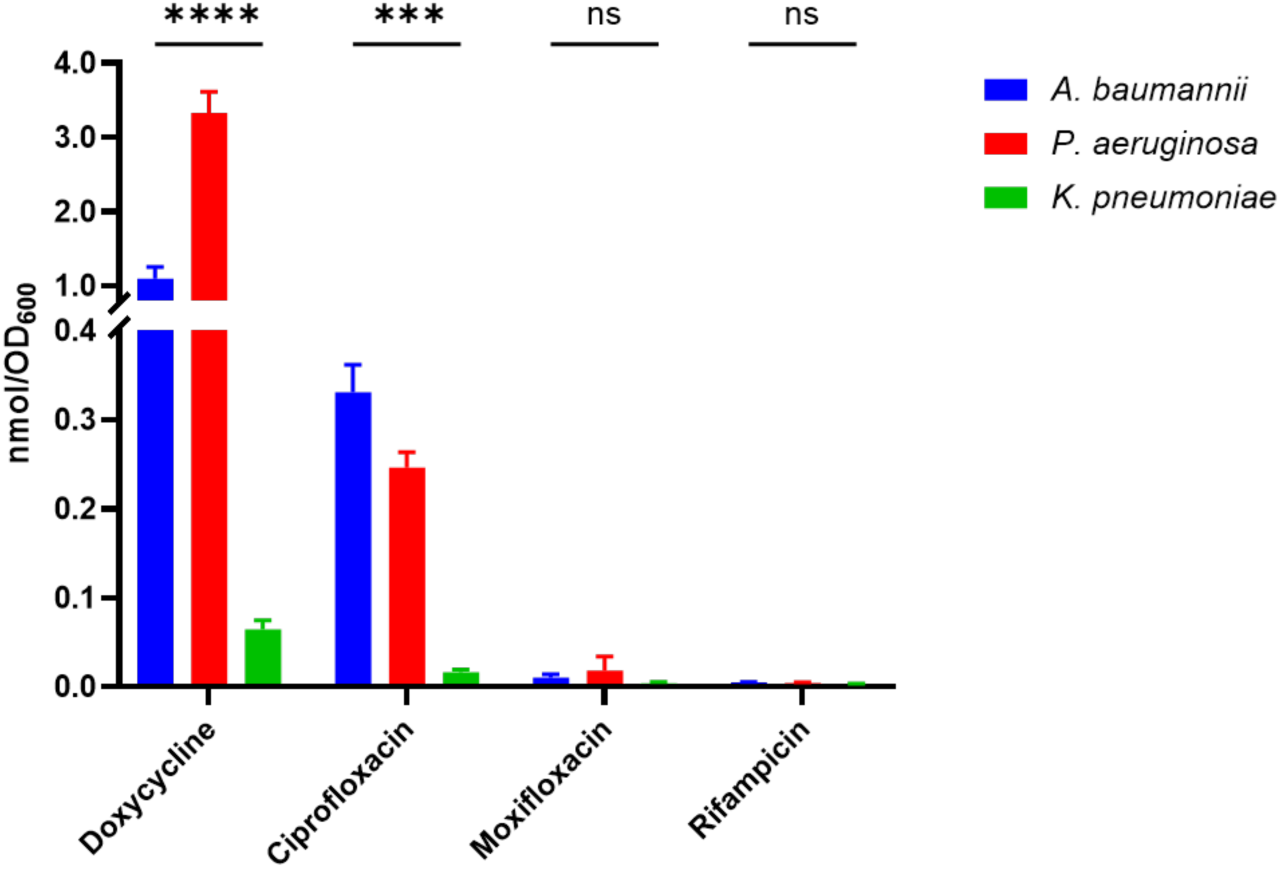
| htIBDM profiling of antibacterial drugs with select ESKPAPE strains. Intrabacterial accumulation of doxycycline, moxifloxacin, ciprofloxacin, and rifampicin (20 µM) in *A. baumannii* (ATCC# 19606), *P. aeruginosa* (ATCC# HER-1018), or *K. pneumoniae* (ATCC# BAA 2146). Data are shown as mean ± SD representative of two independent experiments each conducted in quadruplicate. The amount of accumulated compound as the number of moles was normalized by cell number as approximated by OD600. p-values were determined by two-way ANOVA with Tukey post hoc test for all assays. ns p>0.05, ** p<0.01, *** p<0.001, **** p<0.0001.

### The htIBDM assay characterizes plasmid-mediated multidrug resistance in a clinical isolate of *K. pneumoniae*

To explore the effects of plasmid-mediated resistance in a representative MDR clinical isolate of *K. pneumoniae*, we set out to utilize the htIBDM platform to study drug accumulation in an MDR *K. pneumoniae* strain (70163), its plasmid-cured derivative strain (74189),^39^ and a *qnrB1* knockout strain (75762) derived from 70163 where *qnrB1* is a gene involved in fluoroquinolone resistance with poorly understood function^40^. Using our previously disclosed CRISPR-Cas9 system technology,^41^ curing of the IncF hybrid plasmid (pKPN-K7)^42^ facilitated removal of all resistance genes harbored on it, including *aac(6’)-Ib*-cr^43^ and *qnrB1* (Supplementary Table 3). 70163 still contains known chromosomal mutations (GyrA-83I and ParC-80I) conferring fluoroquinolone resistance^44^ that persist within the plasmid-cured strain 74189. The observation that ciprofloxacin’s intrabacterial accumulation was higher than that for moxifloxacin in previous experiments with the *K. pneumoniae* ATCC# BAA 2146 strain (Fig. 9) and the structural diversity within the fluoroquinolone family of drugs led us to hypothesize that accumulation occupies an important role in resistance to different fluoroquinolones.

Seven fluoroquinolones of clinical relevance were assessed for their MIC versus each of these three strains with doxycycline, rifampicin, and imipenem as control drugs (Table 2). The 70163 strain was not susceptible to any of the seven fluoroquinolones tested. In contrast, the plasmid-cured 74189 strain demonstrated a dramatic (≥8-fold) reduction in the fluoroquinolone MIC values as a likely consequence of losing both the *qnrB1* and *aac(6’)-lb-*cr genes harbored on the plasmid. Furthermore, for all of the fluoroquinolones except norfloxacin, significant differences, of at least fourfold, in the MICs were observed between the plasmid-containing strain 70163 and its derived *qnrB1* knockout strain.

**Table 2.** | MIC values for select antibacterial agents versus MDR *K. pneumoniae* clinical isolate (70163), its plasmid-cured strain (74189), and its *qnrB1* knockout strain (75762). Each MIC value represents the average value from a minimum two independent experiments in µM.

|  | *K. pneumoniae* strain MIC (μM) |  |  |
| --- | --- | --- | --- |
| Drug | 70163 | 74189 | 75762 |
| ciprofloxacin | ≥200 | 6.3 | 50 |
| delafloxacin | 100 | 1.6 | 3.1 |
| gemifloxacin | 50 | 6.3 | 12 |
| levofloxacin | 50 | 6.3 | 12 |
| moxifloxacin | 50 | 6.3 | 12 |
| norfloxacin | >200 | 25 | >200 |
| ofloxacin | 100 | 12 | 12 |
| doxycycline | 50 | 3.1 | 50 |
| rifampicin | 25 | 12 | 25 |
| imipenem | 1.6 | 1.6 | 0.78 |

The fluoroquinolones tested may be generally stratified into two groups with regard to the 74189 strain: high accumulators and low accumulators (Fig. 10). The high accumulators are ciprofloxacin, norfloxacin, and gemifloxacin and the low accumulators are levofloxacin, ofloxacin, moxifloxacin, and delafloxacin. Of the three high accumulators, ciprofloxacin and norfloxacin showed statistically significant decreased accumulation in the MDR plasmid-containing strain (70163) as compared to the plasmid-cured strain (74189), suggesting that decreased intrabacterial accumulation could be a significant contributing factor to the strain’s resistance to those agents. For all of the fluoroquinolones assessed for accumulation in the 75762 versus 70163 strains, we note that the deletion of *qnrB1* fails to result in statistically significant increased drug accumulation. This observation does not support a role for this gene in drug resistance through direct modulation of fluoroquinolone accumulation.

**Fig. 10.**
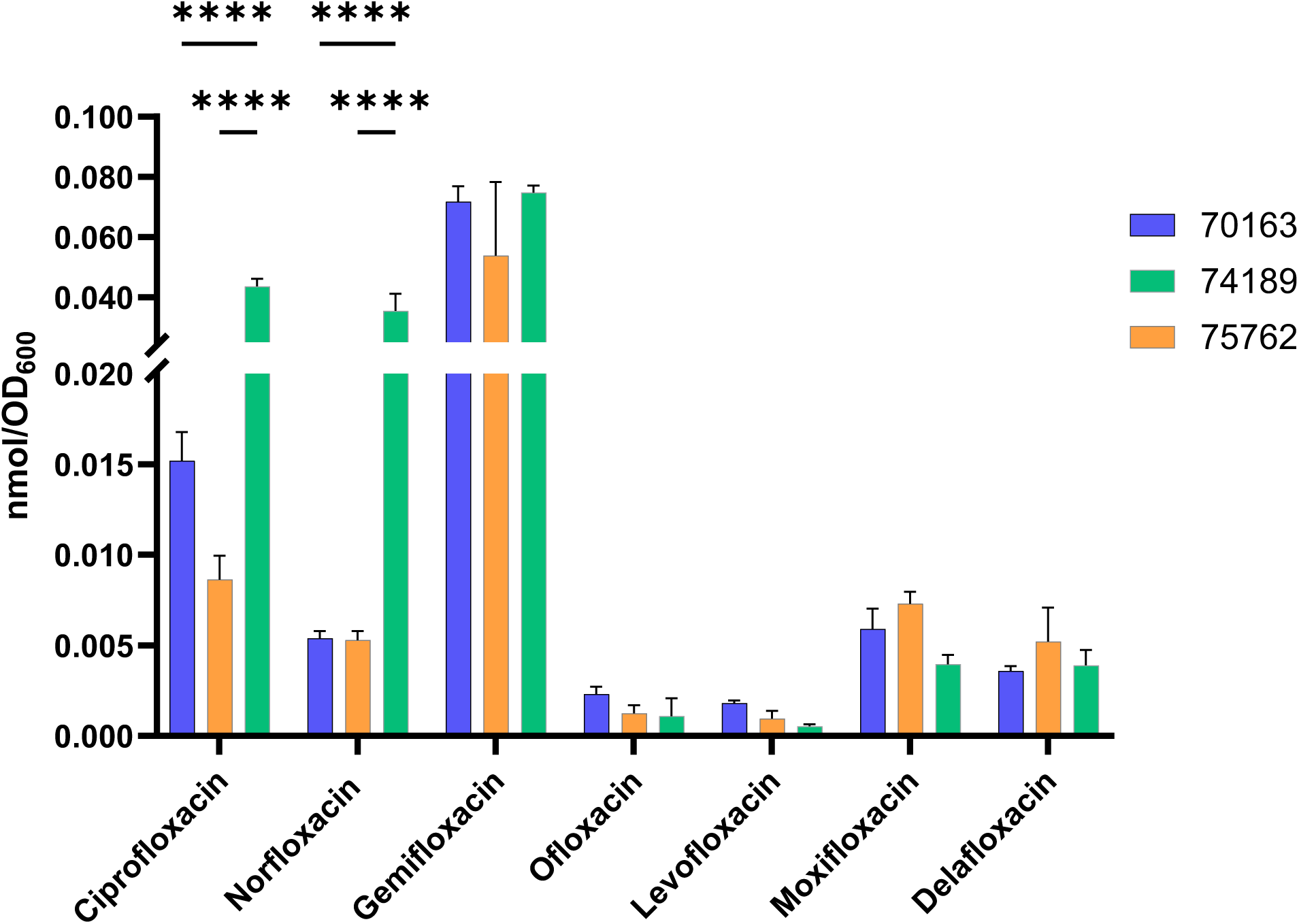
| htIBDM of select antibacterials with a *K. pneumoniae* clinical isolate and its plasmid-cured and *qnrB1* knockout derivative strains. Intrabacterial accumulation of ciprofloxacin, norfloxacin, gemifloxacin, ofloxacin, levofloxacin, moxifloxacin, and delafloxacin (20 µM) in an MDR strain (70163), its plasmid-cured strain (74189) and its *qnrB1* knockout strain (75762). Data are shown as mean ± SD representative of two independent experiments each conducted in quadruplicate. The amount of accumulated compound as the number of moles was normalized by cell number as approximated by OD600. p-values were determined by two-way ANOVA with Šidák post hoc test for all assays. ns p>0.05, ** p<0.01, *** p<0.001, **** p<0.0001. Only statistically significant comparisons amongst accumulations for a given drug are annotated.

## Discussion

Various methods have been previously reported to measure intrabacterial drug accumulation.^45^ Examples include radiometric detection,^46,47^ fluorescence quantification,^48–50^ and LC/MS-based assay.^26,51^ Drug metabolism has not been generally quantified within a bacterium beyond a set of examples from our laboratory and others^21,22,24,52–54^. We assert that an LC/MS-based assay is best suited to provide a direct, label-free quantification of intrabacterial drug accumulation and metabolism while radiometric detection necessitates the synthesis of radiolabeled drug which can be expensive and time-consuming. Furthermore, parent drug and drug metabolites may have the same response profile and may not easily be distinguished. Fluorescence-based measurements are limited by requiring the drug molecule being sufficiently fluorescent^55,56^ or necessitate the addition of a fluorescent moiety that may perturb the parent molecule’s biological activity and/or permeability.

Inspired by the well-established metabolomics platforms originating with Botstein,^57^ Rabinowitz,^57,58^ Rhee,^59^ and a subsequent report from the Dartois laboratory quantifying drug accumulation in *M. tuberculosis*,^60^ we published in 2019 the results of a long-standing effort to design the IBDM platform for *M. tuberculosis*. Our platform represented an amalgamation of the liquid culture approach of Dartois and the sample processing workflows of Botstein, Rabinowitz, and Rhee. Through studies focused on the mechanism of action of two series of nitro-containing heterocyclic antituberculars,^21,22^ we confirmed that our IBDM platform is a useful tool to monitor intrabacterial drug accumulation and metabolism in *M. tuberculosis*. Extensions of the platform were then achieved with *S. aureus*,^24^ uninfected and *M. tuberculosis*-infected J774.1 macrophage-like cells,^20,23^ and Vero cells.^61^ For Gram-negative bacteria, previous reports from other groups have focused on the development of a drug accumulation methodology and we highlight examples from Hergenrother (e.g., *E. coli*^26^ and *P. aeruginosa*^62^), Kern (e.g., *E. coli*^63^), Stover (e.g., *P. aeruginosa*^64^), Tan (e.g., *E. coli*^65^ and *P. aeruginosa*^66^), Zgurskaya (e.g., *E. coli*,^67^ *P. aeruginosa,*^67,68^ and *A. baumannii*^67^) that are label-free in that they do not require a reactive tag on the molecule to quantify its accumulation. In contrast are platforms that require a tag on the analyte drug such as a chloroalkane, a sulfide-linked D-cystine, or azide.^49,50,69^ However, we note the advantage of these approaches to examine localized accumulation (e.g., the cytosol or periplasm). Just as has been shown for the azide-tagged approach,^49^ our IBDM approach can be performed in high-throughput screening format. Validated herein with acceptable Z’ factors for use with *E. coli*, *K. pneumoniae*, *A. baumannii*, and *P. aeruginosa* strains, the htIBDM assay allows the rapid examination of an experimental matrix comprised of a range of both strains and compounds. With this goal in mind, studies have been reported by Widya and colleagues.^70^ This approach, leveraging solid-phase extraction/mass spectrometry, provides excellent throughput while requiring specialized instrumentation that may not be present in the typical academic laboratory. The Zgurskaya laboratory has also applied this technology.^67^ Our htIBDM assay only requires a single-quad LC/MS instrument. Furthermore, we emphasize the facile adaptation of our approach to a high-throughput format that has not yet been reported with the approach of Hergenrother^26^. It should be noted that the methodology used by Hergenrother, adapted from Bazile et al.,^71^ differs from our approach in the utilization of 1) minimal media for the cell culture step in the presence of drug, 2) silicone oil in the cell pelleting step, and 3) CFUs to normalize quantified accumulation.

A primary exemplification of our technology reported herein involves the correlation of measurements of drug accumulation in Gram-negative bacteria with the quantifications of *in vitro* drug efficacy and resistance. We demonstrated with *E. coli* and a small set of antibacterials that changes in drug accumulation inversely correlate with the drug MIC when comparing bacterial strains (e.g., wild type MG1655 versus efflux deficient *ΔtolC*). Furthermore, comparison of the *in vitro* efficacy and accumulation profiles for a set of approved fluoroquinolone drugs with the *K. pneumoniae* MDR clinical strain 70163 and its plasmid-cured derivative strain 74189 demonstrated that removal of the plasmid with known resistance genes can lead to the expected increases in drug susceptibility and drug accumulation. This observation was best demonstrated with ciprofloxacin and norfloxacin; although we note that the trend was not observable for four of the seven approved fluoroquinolones tested. We suspect that their differing accumulation behavior may be explained with further contemplation of the effect of their varying chemical substitutions of the fluoroquinolone scaffold. This knowledge, coupled with an understanding of their respective intracellular target engagement/s, should be useful in developing a better understanding of their drug resistance profiles. These studies add to a set of examples in the literature led by the Zgurskaya laboratory where 1) differences in compound growth inhibition as quantified by MIC have been compared to direct measurements of drug accumulation with a single bacterial strain^66,67,72–75^ and 2) quantitative changes in drug accumulation were correlated with mutations that affect either the outer membrane or efflux,^67,68,72,76^ porins,^67,68^ or transporters^77^. Interestingly, we demonstrated that the CRISPRi knockdown of *qnrB1* affects the MIC values of the fluoroquinolone class without significantly perturbing accumulation. This outcome is consistent with the lack of data linking this gene to fluoroquinolone accumulation.^40^ These results with *K. pneumoniae* underscore the potential of our approach to illuminate the role of Gram-negative genes in drug resistance as viewed through the lens of drug accumulation. In particular, we see the promise of combining high-throughput genetic approaches such as CRISPRi^78,79^ with our htIBDM platform.

Another application of our technology is in the realm of drug combination studies. We have probed the interaction between rifampicin and indacaterol, inspired by the elegant study of Blainey leveraging a high-throughput nanoliter scale screen^31^. In a dose-dependent fashion, we demonstrated the ability of indacaterol to augment the accumulation of rifampicin within *E. coli* and effectively increase the potency of rifampicin. While molecules such as colistin^26^ and pentamidine^80^ have been demonstrated to sensitize Gram-negative bacteria to typical Gram-positive selective agents, we are only aware of the study by Hergenrother that showed colistin to quantitatively augment the accumulation of novobiocin within the bacterium^26^. We also note studies on compound accumulation in *E. coli* in the presence of efflux inhibitors^65^. It is our expectation that further studies of the accumulation and metabolism dynamics of drug combination systems will be illuminated through the methods as outlined herein. It is intriguing to consider how traditional checkerboard studies used to infer interactions between two compounds, or their higher dimensional related approaches,^81^ could benefit from consideration of how combinations of drugs influence their individual accumulation within cells. Such knowledge gained from these studies should be critical to the rational development of combination therapies for diseases necessitating multi-drug therapy.

Returning to the quantification of intrabacterial drug accumulation in single-compound studies, we speculate the field will be able to further stratify the results from high-throughput growth inhibition studies through the identification of compounds that do or do not achieve significant accumulation within the targeted cell. The generated screening data would augment currently available datasets for the construction and validation of machine learning models for drug accumulation and metabolism. Such computational approaches have been undertaken for *P. aeruginosa*^68,82,83^ and *E. coli*^26^. Computational predictions and the actual experimental determinations of drug accumulation should bolster antibacterial medicinal chemistry optimization to further dimensionalize and inform structure-activity relationship (SAR) studies. Early examples of this approach may be found in the literature,^66,73,74^ although they are far from commonplace.

Given our previous work quantifying drug accumulation and metabolism within *M. tuberculosis*^21,22^ and *S. aureus*^24^, we were interested in investigating the significance of drug metabolism within Gram-negative bacteria. We focused on *E. coli* and specifically the MG1655 strain. Rifampicin was chosen to probe intrabacterial biotransformations due to the commercial availability of rifampicin N-7-oxide^84,85^ metabolite (Supplementary Table 4) and the extensive optimization efforts with our IBDM protocol performed with the parent compound rifampicin. Initial studies were conducted at 20 μM rifampicin, and sample collection was performed at 10 and 60 min. The metabolite was not detected under these conditions. However, when 200 µM rifampicin was added to a culture OD600 of 0.77, we observed the formation of the N-7-oxide metabolite at t = 90 min (Supplementary Fig. 10). Interestingly, the MIC of the rifampicin N-7- oxide is ≥16-fold higher than that for the parent compound rifampicin (Supplementary Table 5), highlighting the potential for deactivating intrabacterial drug metabolism^52^. Further studies are needed to characterize the formation of this metabolite and others in Gram-negative bacteria.

Limitations to the methodology have already been discussed in detailed review of a somewhat similar protocol by Hergenrother, differing mainly in sample processing as discussed above^86^. These limitations are related to the quantification of compound accumulation with an LC/MS based methodology, the potential for compound covalent attachment to intracellular proteins, and an inability to provide information as to compound localization within the cell.

In conclusion, we reflect on the 2020 discussion of drug penetration in Gram-negative bacteria by Zgurskaya and Tan and their concluding remarks on remaining challenges^87^. Firstly, we assert the IBDM platform and the results discussed herein address the need for a high-throughput drug accumulation platform that is readily available to most laboratories. Secondly, our studies of drug resistance mutations in laboratory and clinical strains complement the authors’ publications in laying further groundwork for understanding how bacterial genes perturb drug accumulation and, hence, drug efficacy/resistance. These and other challenges in chemical biology and drug discovery as they pertain to small molecule-microbe interactions are open to being addressed through our platform in intrabacterial drug accumulation.

## Methods

### Commercially sourced bacterial strains and reagents

*E. coli* K-12 MG1655 ^88^ and the isogenic efflux deficient strain Δ*tolC* ^89,90^ were obtained from the Brynildsen lab (Princeton University). *A. baumannii* (ATCC# 19606)*, P. aeruginosa* (ATCC# HER-1018), or *K. pneumoniae* (ATCC# BAA 2146) were sourced from the ATCC. *K. pneumoniae* strain 70163 was accessed from the Hackensack Meridian Health Center for Discovery & Innovation clinical collection. Small molecules were sourced as follows: Chem-Impex – doxycycline, moxifloxacin, ciprofloxacin, and ofloxacin; MedChemExpress – indacaterol; TCI America – rifampicin; AA Blocks – gemifloxacin and delafloxacin; Activate Scientific – norfloxacin and levofloxacin; LGC Standards – rifampicin- d8, rifampicin N-7-oxide; – verapamil.

### Construction of *K. pneumoniae* strain 74189

To eliminate the IncF hybrid plasmid pKPN-K7 from strain 70163, a targeted plasmid curing approach was employed using the conjugative CRISPR-Cas9 vector pLCasCureT, as described by Yen and co-workers^41^. A previously constructed version of this vector, encoding a guide RNA specific to the IncF replicon, was conjugated into the parental strain via mating. Following conjugation, expression of Cas9 was induced by the addition of arabinose to activate plasmid targeting. Transconjugants were then selected and screened for plasmid loss. Successful curing of pKPN-K7 was confirmed by the absence of IncF replicon-specific PCR amplicons.

### Construction of *K. pneumoniae* strain 75762

A twenty nucleotide guide RNA (N20) adjacent to a PAM site within the *qnrB1* locus was designed and cloned into the pSGKP plasmid, following the method of Wang and colleagues^91^. The parental *K. pneumoniae* strain 70163 was co-transformed with three components: the pCasKP plasmid encoding the Cas9 nuclease and the lambda red recombination system, the pSGKP plasmid carrying the guide RNA, and a separate double-stranded DNA repair template containing homologous flanking sequences. This approach enabled the precise deletion of *qnrB1*. Gene deletion was confirmed by Sanger sequencing.

### Alamar blue assay to determine MIC of drugs against bacterial strains

A broth microdilution assay was performed to determine the MIC of drugs tested against strains of *E. coli*, *A. baumannii*, *K. pneumoniae*, and *P. aeruginosa*. Assays were performed in two biological replicates and the MIC of each drug was determined in duplicate wells. Control wells contained DMSO. An overnight culture of bacteria was subcultured 1:100 in fresh LB media and grown at 37 °C and 180 rpm until reaching an OD600 of 0.2 – 0.4. The bacterial culture was then diluted 1:1000 in LB media and added to a 96-well microtiter plate containing different concentrations of serially diluted drug (final volume 100 µL). The plate was grown at 37 °C without shaking. After 24 h, the MIC was determined using either visual inspection of the cell pellet or via the addition of Alamar blue (Invitrogen). In the case of the later, 10% Alamar blue was added to the cultures and returned to the incubator for 3 h. The MIC was determined as the lowest concentration of drug resulting in complete growth inhibition by the chosen method of observation.

### Intrabacterial drug metabolism assay (IBDM)

The IBDM assay was performed with the appropriate Gram-negative bacterial strain with select drugs. All solutions added to cell pellets were pre-chilled over ice and all centrifugations were performed at 4 °C to minimize further metabolic processes post-sample collection. An overnight culture of bacteria was subcultured 1:100 in fresh LB media and grown at 37 °C and 180 rpm until mid-log phase (OD600 0.5 – 0.6). A DMSO treated culture was used as the control. At the appropriate timepoint (e.g., 10, 30, and/or 60 min) at least three 1 mL samples were collected and centrifuged at 7,500 rpm for 5 min. The cells were then washed with ice cold 0.85% NaCl solution. The resulting cell pellet was quenched with an ice-cold mixture of 2:2:1 CH3OH:CH3CN:H2O (MAW). The resulting solution was then lysed by four cycles of freeze-thaw (5 min freeze/30 s thaw) in a dry ice/acetone bath. The cell lysate was subsequently centrifuged at 13,000 rpm for 5 min. 700 µL of supernatant were then transferred to a 0.22-micron filter tube and centrifuged at 13,000 rpm for 10 min to filter cellular debris. Samples were stored in a -80 °C freezer until LC/MS analysis.

### LC/MS analysis of biological lysates

Accumulation was measured using an Agilent Infinity II liquid chromatography system coupled with an Agilent 6125 single quadrupole mass spectrometer. Liquid chromatography separation was achieved using an Agilent Poroshell 120 C-18 column with 1.9 µm particle size, 2.1 mm internal diameter, and 50 mm length. The solvent system consisted of CH3CN and H2O supplemented with 0.1% formic acid. The gradient involved four main phases: 1) an isocratic phase from 0 – 0.3 min at 5% CH3CN, followed by 2) 0.3 – 0.6 min increasing CH3CN to 30%, 3) 0.6 – 3.3 min increasing CH3CN from 30% to 95%, and 4) an isocratic phase from 3.3 – 3.6 min at 95% CH3CN. This was followed by a post-run for 1 min at 95% H2O. The flow rate was set to 0.5 mL/min, and the column temperature was set to 40 °C. These parameters were held constant for all drugs tested. For mass spectrometry, all analyses were performed using an API-ES ion source in positive polarity. The following spray chamber parameters were used: nebulizer pressure of 40 psi, capillary voltage of 3000 V, drying gas temperature of 350 °C, and drying gas flow of 12.0 L/min. Two main methods were used to quantify intrabacterial accumulation. Firstly, a scan method ranging from m/z 100 – 1000 was used to investigate the retention time of the drug being tested and its ionization. Then a selective ion monitoring (SIM) method was employed to maximize sensitivity. Each SIM method was set up to search for two target masses: the drug tested and the internal standard. For example, the SIM method for rifampicin searched for m/z 823 for rifampicin and m/z 831for rifampicin-d8. For all other drugs, verapamil was used as the internal standard (m/z 455).

### Calculation of drug accumulation from ion counts

To quantify drug accumulation expressed as nmols/OD600, drug calibration curve samples were prepared and submitted for LC/MS analysis on the same day the biological samples were run on the instrument to control for day-to-day variations in instrument sensitivity. 10 – 15 serial dilutions (typically, 10 – 0.00061 µM) of drug were prepared in 250 µL of 2:2:1 CH3CN/MeOH/H2O. 5.0 µL from a stock solution of internal standard (typically ranging from 10 – 100 µM) was added to all calibration curve samples. Peak areas obtained from LC/MS analysis were then plotted using Microsoft Excel 365 and the slope of the best fitting regression line (m) was used in equation 1 to calculate nmols of drug from peak area obtained from biological samples.

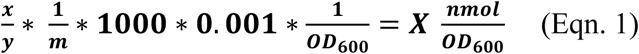

Equation 1 calculates X as the drug accumulation expressed in nmol/OD600 , where “x” and “y” represent the peak areas of drug and internal standard, respectively. “m” represents the slope of the regression line obtained from the calibration curve. Since the ion count generated by the LC/MS is a unitless measure, x/y ratio was divided by “m” to afford a number µM as its units, which was then multiplied by 1000 to convert it to nM. Then, the number was multiplied by 0.001 (to account for the 1 mL sample volume) to yield a final number with nmol units. The final step was to normalize nmols by OD600 of the bacterial culture at each time point to yield a final number with units nmol/OD600. Comparison of the calibration curve slope determined for each drug in either 2:2:1 CH3CN/MeOH/H2O or the lysate from *E. coli* MG1655 grown to OD600 ∼ 0.5 – 0.6 showed a negligible matrix effect (Supplementary Table 6). Thus, when conducting single-compound and high-throughput IBDM experiments with Gram-negative bacteria other than *E. coli*, we determined the calibration curve slope using a 2:2:1 CH3CN/MeOH/H2O solution of the drug.

### Identification of rifampicin metabolites using an LC-TOF system

To search for metabolites of rifampicin, an Agilent 1260 Infinity II liquid chromatography, fitted with an InfinityLab Poroshell 120 EC-C18 column (2.1 x 100 mm, 2.7 µm) and coupled with an Agilent 6230 time-of-flight (TOF) mass spectrometer, was used to analyze the IBDM samples. LC and spray chamber parameters were identical to the LC/MS conditions listed above with the exception of a longer run time (22 min) to accommodate for the lower flow rate (0.3 mL/min) used in the 1260 Infinity II LC system. The Agilent Technologies MassHunter Quant software version 10.0 combined with the Agilent Personal Compound Database and Library (PCDL version 8.0) were used to search for the chosen metabolite (Supplementary Table 4) in the rifampicin treated sample (200 µM) and the DMSO treated control. Candidate metabolites were first identified as transformations of the parent compound that were observed in drug-treated samples and not in the DMSO-treated control. Secondly, they were confirmed by matching their m/z and retention time to the commercially obtained standard.

### Checkerboard assay for assessing synergy of rifampicin with indacaterol

A checkerboard assay, as previously described,^92^ assessed interactions between rifampicin and indacaterol with the *E. coli* MG1655 strain. An overnight culture of bacteria was subcultured 1:100 in LB media and incubated at 37 °C and 180 rpm until an OD600 of 0.5 was reached. The bacterial culture was then diluted 1:1000 in LB media and added to the wells in a 96-well microtiter plate containing indacaterol (0 – 800 µM) and rifampicin (0 – 50 µM) concentrations in serial dilutions. The plate was then incubated without agitation at 37 °C for 24 h. 10% Alamar blue (Invitrogen) was then added to the plate which was returned to the incubator for 3 h. The MIC was determined as the lowest concentration of drug resulting in complete growth inhibition by visual inspection. Drug interactions were determined by calculating fractional inhibitor index (FICI) using equation 2, where “A” represents MIC of drug A in combination and “B” represents MIC of drug B in combination (FIC ≤ 0.5, synergy; 0.5 < FIC < 4.0, neither synergy nor antagonism; FIC ≥ 4.0, antagonism).^34^

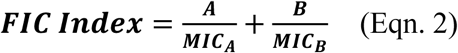

### Cell Staining

Changes in *E. coli* outer membrane permeability as modulated by indacaterol and/or rifampicin were quantified by flow cytometry using TO-PRO-3 (Invitrogen). A culture of *E. coli* MG1655 was diluted 1:100 in LB media and grown at 37 °C with shaking (180 rpm) until an OD600 of 0.50 – 0.60 was reached. To the cells was then added DMSO (1% in LB media) as a negative control, 2.5 µM polymyxin B as a positive control, or the tested concentrations of rifampicin and/or indacaterol. To maintain consistency with our IBDM studies, we incubated the bacterial cells with drugs or control for 1 h. Cells were then pelleted at 3,000 rpm for 10 min at 4 °C and then washed with PBS twice. The supernatant was discarded, and the cells were diluted in fresh PBS to afford an OD600 = 0.01 (ca. 10^6^ CFU/mL)^93^. TO-PRO-3 (1.0 µM) was added to the cells, and the cells were incubated with the dye at rt in the dark for 10 min.

### Flow Cytometry

Flow cytometry assays were conducted with the BD LSRII (BD Biosciences). TO-PRO 3 was detected using APC with an excitation of 633 nm and a 660/20 bandpass filter. Results were analyzed using BD FlowJo software (version 10.7.2, BD Biosciences).

### High-throughput interbacterial drug metabolism (htIBDM) assay

Initially Z’ ^36^ was determined as follows. An *E. coli* MG1655 culture was grown in LB media at 37 °C with shaking at 180 rpm to an OD600 of 0.55 – 0.65. 1.1 mL of the culture was then transferred to each well of a 96-well deep well plate (Agilent # 201240-100) and sealed with a seal mat (Agilent #201158-100). 10 µM of doxycycline was added to all odd-numbered wells, and DMSO was added to all even-numbered wells. The plate was then incubated at 37 °C with shaking at 280 rpm for 1 h. 100 µL from each well in the original plate was transferred to a separate 96-well plate to obtain OD600 measurements using an Agilent BioTek Synergy HTX Multimode Reader. The original plate was then centrifuged at 4,000 rpm and 4 °C for 10 min, washed twice with ice-cold 0.85% aqueous NaCl, and quenched with ice-cold 2:2:1 CH3CN/CH3OH/H2O. Bacterial cells were then lysed by one cycle of freeze-thaw and 3 x 5 min cycles of sonication followed by centrifugation for 10 min at 4,000 rpm. The cell lysate (250 µL) was subsequently filtered using a filter plate (Agilent #203980-100), and flow-through was collected in a collection plate (Nunc #260251). The resulting filtrate was used for LC/MS analysis as described above. The same protocol that was used for the Z’ calculation was used for htIBDM studies with the *E. coli*, *A. baumannii*, *P. aeruginosa*, and *K. pneumoniae* strains tested with the exception that drugs were incubated with each strain in quadruplicate at 20 µM.

## Statistical analyses

All IBDM assays were performed in at least triplicate with two biological replicates. All statistical analysis was performed using GraphPad Prism (version 10.0.0) and are detailed in the caption for each figure.

## Data availability

The datasets generated during and/or analyzed during the current study are available in the Supplementary Information.

## Funding

NIH U19AI142731, U19AI109713.

## Acknowledgements

Dr. Sukhwinder Singh (Rutgers University) for assistance with the flow cytometry measurements and analysis. Professors Paul C. Blainey (MIT), Mark P. Brynildsen (Princeton University), Emily R. Derbyshire (Duke University), and M. Sloan Siegrist (UMass-Amherst) in addition to Marcella Widya (UCB) for helpful discussions.

## Author contributions

A.G., S., Y.-M.A., and J.S.F. conceptualized the project. A.G., S., Y.-M.A., B.N.K., and J.S.F. developed the methodology. Y.-M.A. and P.R.B. performed initial validation.

A.G. and S. conducted formal analyses. A.G., S., A.B., K.S., and A.J.T. conducted investigations.

A.G, S., and J.S.F. wrote the original draft. A.G., S., A.J.T., B.N.K., and J.S.F. reviewed and edited the manuscript. B.N.K. and J.S.F. supervised the project. J.S.F. acquired funding.

## Competing interests

The authors declare no competing interests.

## Notes

### Competing Interest Statement

The authors have declared no competing interest.

