## Supplemental Information for "The Quantification of Drug Accumulation within Gram-Negative Bacteria"

**Table of Contents**

Figure S1. Accumulation of rifampicin in *E. coli* MG1655 in three different culture volumes. Page S4

Figure S2. Accumulation of rifampicin in *E. coli* MG1655 at four different time points. Page S5

Figure S3. Doxycycline and rifampicin accumulation in *E. coli* MG1655 normalized by $OD_{600}$ versus colony-forming units (CFUs). Page S6

Figure S4. Moxifloxacin, doxycycline, and rifampicin time-dependent accumulation in *E. coli* MG1655. Page S7

Figure S5. Moxifloxacin, doxycycline, and rifampicin dose-dependent accumulation in *E. coli* MG1655 at t = 60 min. Page S8

Figure S6. Checkerboard assay showing interaction between rifampicin and indacaterol with respect to the *E. coli* MG1655 strain. Page S9

Figure S7. DMSO treated *E. coli* MG1655 cells in the presence of TO-PRO-3 dye. Page S10

Figure S8. Single compound IBDM of rifampicin, moxifloxacin, novobiocin, ciprofloxacin and doxycycline in *E. coli* MG1655. Page S11

Figure S9. Effect of number of washes on the accumulation of novobiocin in *E. coli* MG1655. Page S12

Figure S10. Chromatograms of rifampicin and its N-7-oxide metabolite metabolite in cell lysate compared to a commercial standard. Page S13

Table S1. MIC of rifampicin against select Gram-negative ESKAPE strains. Page S14

Table S2. Molecular weight (MW; g/mol) and h_logD values for compounds tested. Page S15

Table S3. Genotypes of clinical isolate 70163 and its plasmid-cured 74189 and *qnrB1* knockout 75762 derivative strains. Page S16

Table S4. Chemical formulas and exact masses for rifampicin and its investigated metabolite. Page S17

Table S5. MICs of rifampicin and rifampicin N-7-oxide against *E. coli* MG1655. Page S18

Table S6. Matrix effects for moxifloxacin, doxycycline, and rifampicin. Page S19

It should be noted that all data for the IBDM runs discussed in this publication may be found in the file: Supplementary_File_1.XLSX

**Supplementary Figure 1. Accumulation of rifampicin in *E. coli* MG1655 in three different culture volumes.** Data are shown as mean ± SD of 2 – 3 technical replicates of rifampicin (Rif;10 μM) incubated with bacteria with three different culture volumes (15, 25, and 125 mL). A Kruskal-Wallis test with Dunn post-hoc test was performed, and no statistically significant difference was found between rifampicin accumulation in all three culture volumes ($p > 0.05$).

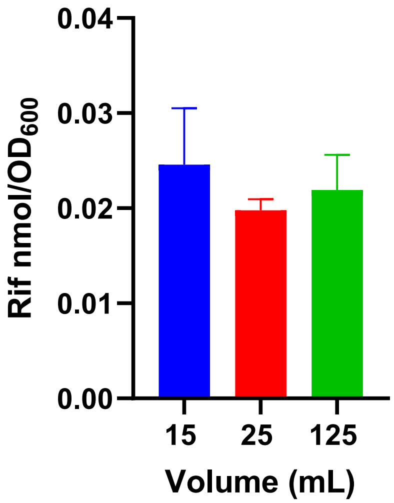

**Supplementary Figure 2. Accumulation of rifampicin in *E. coli* MG1655 at four different time points.** Data are shown as mean ± SD of 2 – 3 technical replicates for rifampicin (Rif; 10 μM) incubated with a 25 mL culture of *E. coli* MG1655. A Kruskal-Wallis test with Dunn post-hoc test was performed, and no statistically significant difference was found between rifampicin accumulation at all time points ($p > 0.05$).

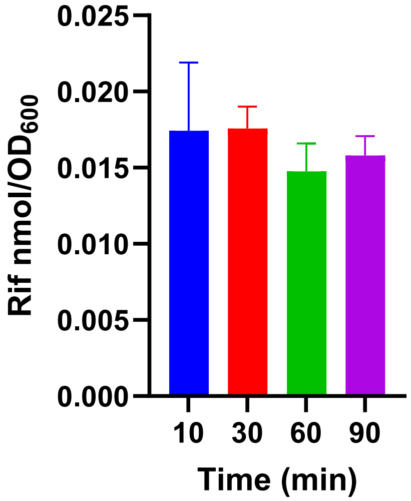

**Supplementary Figure 3. Doxycycline and rifampicin accumulation in *E. coli* MG1655 normalized by $OD_{600}$ versus colony-forming units (CFUs).** Doxycycline and rifampicin accumulation at 10 μM compound, normalized by a) $log_{10}$ CFUs compared to b) $OD_{600}$. Data are shown as mean ± SD representative of two independent experiments each conducted in quadruplicate. p-values were determined by two-way ANOVA with Šídák post hoc test. ns p>0.05, ** p<0.01, *** p<0.001, **** p<0.0001.

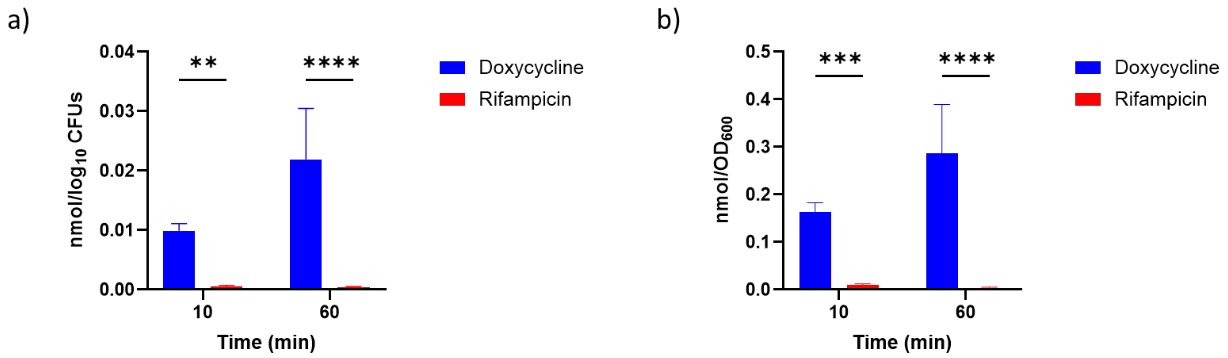

**Supplementary Figure 4. Moxifloxacin, doxycycline, and rifampicin time-dependent accumulation in *E. coli* MG1655.** a. Moxifloxacin time-dependent accumulation at 2.5 and 5.0 μM. b. Doxycycline time-dependent accumulation at 7.5 and 15 μM. c. Rifampicin time-dependent accumulation at 10 and 20 μM. Data are shown as mean ± SD for a representative of two independent experiments each conducted in triplicate. The amount of accumulated compound as the number of moles was normalized by cell number as approximated by $OD_{600}$. p-values were determined by two-way ANOVA with Tukey post hoc test. ns p>0.05, ** p<0.01, *** p<0.001, **** p<0.0001. For each drug at each of the two concentrations assessed, the following pairwise comparisons demonstrated ns p>0.05 : 10 min vs. 30 min, 10 min vs. 60 min, and 30 min vs. 60 min.

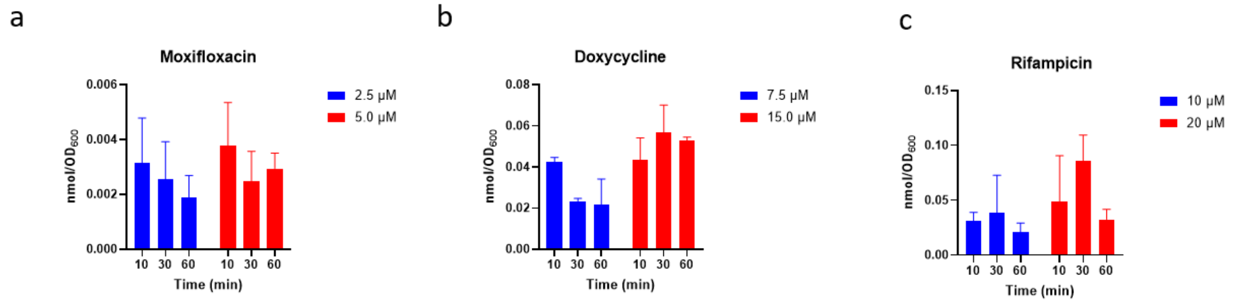

**Supplementary Figure 5. Moxifloxacin, doxycycline, and rifampicin dose-dependent accumulation in *E. coli* MG1655 at t = 60 min.** Data are shown as mean ± SD for a representative of two independent experiments each conducted in triplicate for a) moxifloxacin, b) doxycycline, and c) rifampicin. The amount of accumulated compound as the number of moles was normalized by cell number as approximated by $OD_{600}$. p-values were determined by unpaired t-tests comparing the two concentrations for each drug. ns p>0.05, ** p<0.01, *** p<0.001, **** p<0.0001.

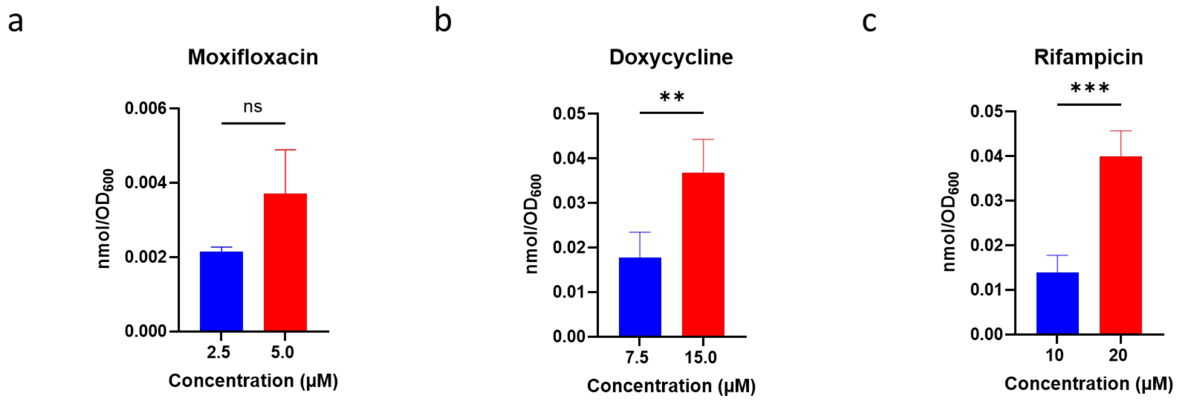

**Supplementary Figure 6. Checkerboard assay showing interaction between rifampicin and indacaterol with respect to the *E. coli* MG1655 strain.** Alamar blue was used to determine the MIC of each drug as the lowest concentration with no cell growth by visual inspection. Drug interactions were determined by calculation of the fractional inhibitor (FIC) index (FIC ≤ 0.5, synergy; 0.5 < FIC < 4.0, neither synergy nor antagonism; FIC ≥ 4.0, antagonism). The rifampicin MIC in combination was ≤0.78 µM, the indacaterol MIC in combination was 50 µM, and, thus, the FIC index was calculated to be ≤0.19.

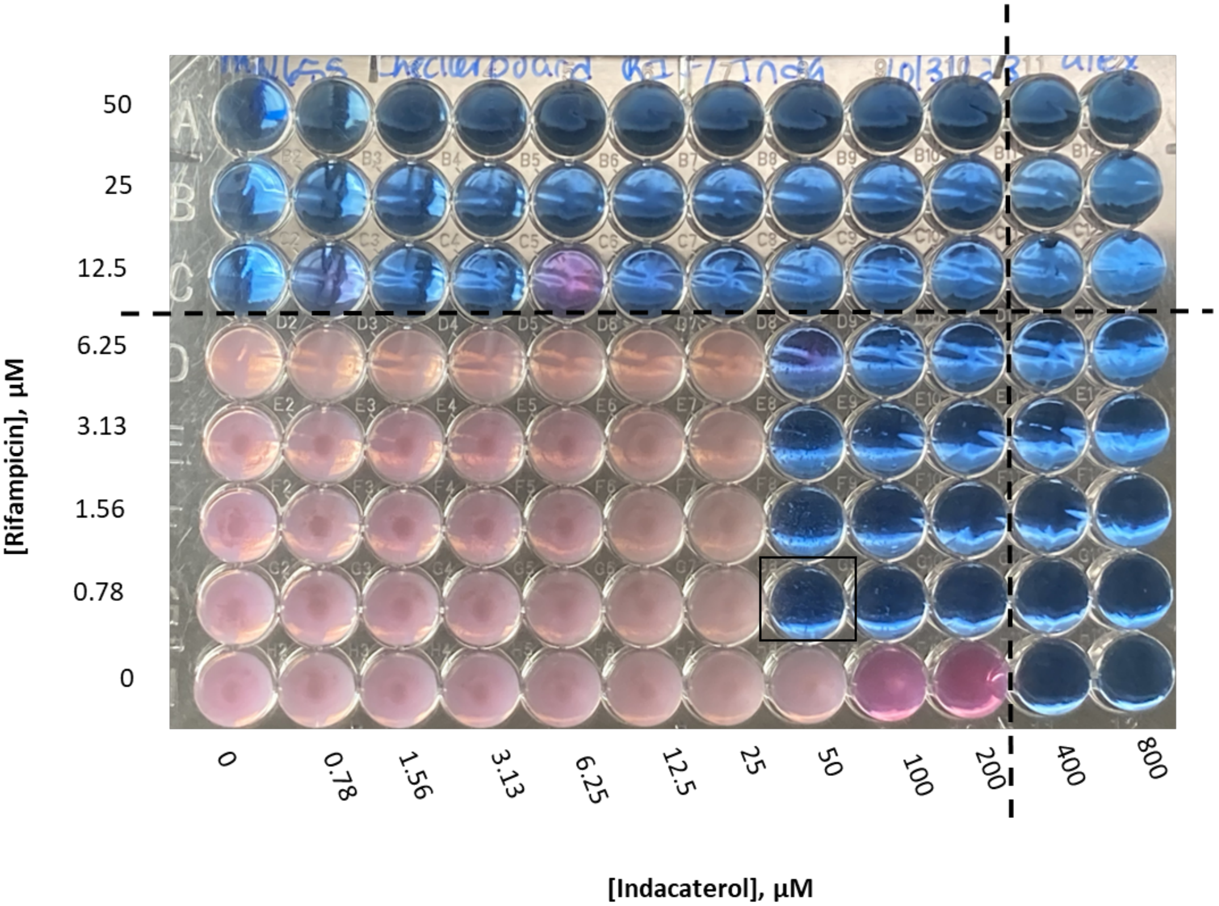

**Supplementary Figure 7. DMSO treated *E. coli* MG1655 cells in the presence of TO-PRO-3 dye.** These data represent the negative control for dye uptake where the percent of cells that took up the dye is shown. The y-axis represents the forward scatter area and the x-axis represents the fluorescence of TO-PRO-3. Results are representative of three independent experiments.

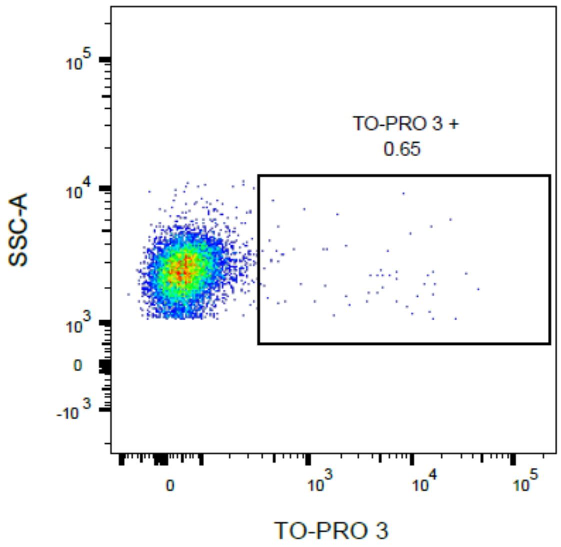

**Supplementary Figure 8: Single compound IBDM assay accumulation of rifampicin, moxifloxacin, novobiocin, ciprofloxacin and doxycycline in *E. coli* MG1655.** Data are shown as mean ± SD representative of two independent experiments each conducted in quadruplicate for the drug concentration of 20 µM. The amount of accumulated compound as the number of moles was normalized by cell number as approximated by $OD_{600}$. p-values were determined by a one-way ANOVA with Tukey post hoc test. ns p>0.05, ** p<0.01, *** p<0.001, **** p<0.0001.

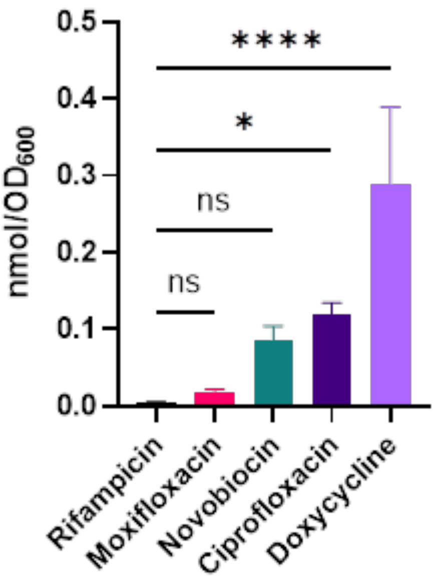

**Supplementary Figure 9. Effect of number of washes on the accumulation of novobiocin in *E. coli* MG1655.** Data are shown as mean ± SD conducted in quadruplicate. The amount of accumulated compound as the number of moles was normalized by cell number as approximated by $OD_{600}$. p-values were determined by a Kruskal-Wallis test with Dunn post hoc test. ns p>0.05, ** p<0.01, *** p<0.001, **** p<0.0001.

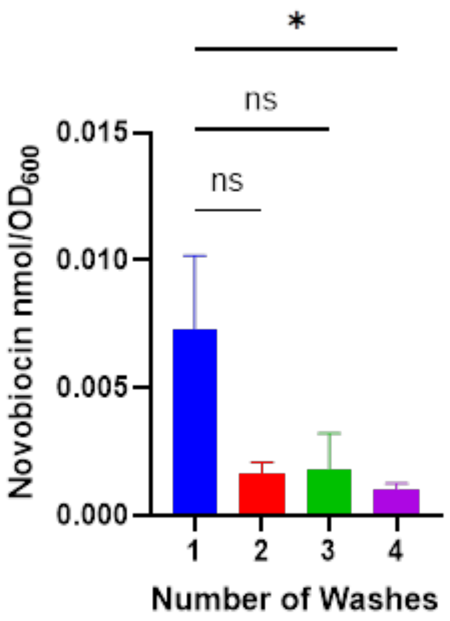

**Supplementary Figure 10. Chromatograms of rifampicin and its N-7-oxide metabolite in cell lysate compared to a commercial standard.** Chromatograms of rifampicin N-7-oxide authentic standard (red), rifampicin N-7-oxide detected in *E. coli* MG1655 (green) and the co-injection of the biological sample and rifampicin N-7-oxide authentic standard (orange).

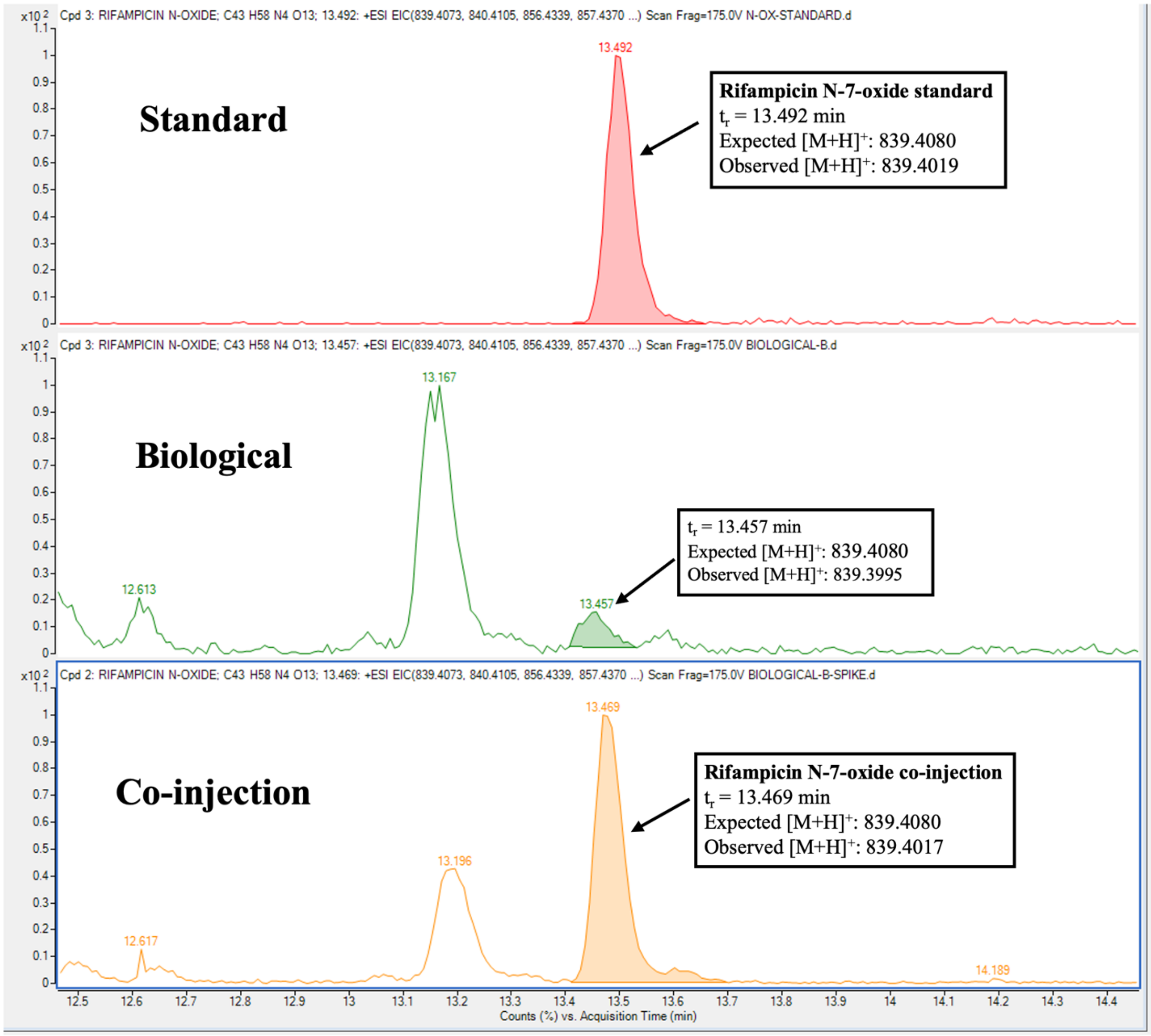

**Supplementary Table 1. MIC of rifampicin against select Gram-negative ESKAPE strains.** MIC values were calculated as the mean of two independent experiments, each performed in duplicate.

| **Bacterial Strain (ATCC #)** | **Rifampicin MIC (µM)** |
|---|---|
| *P. aeruginosa* (HER-1018) | 12 |
| *K. pneumoniae* (BAA 2146) | 25 |
| *A. baumannii* (ATCC#19606) | 0.78 |

**Supplementary Table 2. Molecular weight (MW; g/mol) and h_logD values for compounds tested.** Descriptors were generated by the Molecular Operating Environment (MOE) software package 2024.06.

| Drug | MW (g/mol) | h_logD (pH 7) |
|---|---|---|
| novobiocin | 612.6 | 3.4 |
| rifampicin | 822.9 | 2.2 |
| doxycycline | 444.4 | -0.073 |
| ciprofloxacin | 331.3 | -0.3 |
| moxifloxacin | 401.4 | -1.1 |

**Supplementary Table 3. Genotypes of clinical isolate 70163 and its plasmid-cured 74189 and *qnrB1* knockout 75762 derivative strains.** Curing of the IncF hybrid plasmid (pKPN-K7-1) facilitated removal of all resistance genes harbored on it. 70163 contains chromosomal mutations conferring fluoroquinolone resistance that persist within the plasmid-cured strain 74189. 75762 is the *qnrB1* knockout of the 70163 strain.

| Strain | Sequence Type (ST) | Plasmid Content | Plasmid Encoded Aminoglycoside Resistance | Plasmid Encoded Fluoroquinolone Resistance | Chromosomal Resistance Mutations |
|---|---|---|---|---|---|
| 70163 | ST307 | IncF: pKPN-K7-1 | *aac(6')-Ib-cr* | *qnrB1* | GyrA-83I; ParC-80I |
| 74189 | ST307 | Plasmid-free | | | GyrA-83I; ParC-80I |
| 75762 | ST307 | IncF: pKPN-K7-1 | *aac(6')-Ib-cr* | | GyrA-83I; ParC-80I |

**Supplementary Table 4. Chemical formulas and exact masses for rifampicin and its investigated metabolite.**

| Molecule | Chemical Formula | Exact Mass (amu) | Expected $[M+H]^+$ |
|---|---|---|---|
| rifampicin | $C_{43}H_{58}N_4O_{12}$ | 822.4051 | 823.4131 |
| rifampicin N-7-oxide | $C_{43}H_{58}N_4O_{13}$ | 838.4000 | 839.4080 |

**Supplementary Table 5. MICs of rifampicin and rifampicin N-7-oxide against *E. coli* MG1655.** Each MIC value represents the average value from a minimum of two independent experiments.

| **Compound** | **MIC against *E. coli* MG1655 (μM)** |
|---|---|
| rifampicin | 12 |
| rifampicin N-7-oxide | >200 |

**Supplementary Table 6. Matrix effects for moxifloxacin, doxycycline, and rifampicin.** Calibration curves were constructed in $CH_3OH$:$CH_3CN$:$H_2O$ (2:2:1 MAW) and in *E. coli* MG1655 biological lysate in MAW. The matrix effect was quantified by dividing the slope of the calibration curve (sensitivity) in biological lysate by the slope of the calibration curve in MAW. Values are expressed as the mean from three experiments.

| Drug | Sensitivity in MAW | Sensitivity in Biological Lysate | Matrix Effect |
|---|---|---|---|
| moxifloxacin | 0.67 | 0.55 | 0.82 |
| doxycycline | 0.35 | 0.36 | 1.0 |
| rifampicin | 0.34 | 0.43 | 1.3 |
